# A chromosome scale assembly of the model desiccation tolerant grass *Oropetium thomaeum*

**DOI:** 10.1101/378943

**Authors:** Robert VanBuren, Ching Man Wai, Jens Keilwagen, Jeremy Pardo

## Abstract

*Oropetium thomaeum* is an emerging model for desiccation tolerance and genome size evolution in grasses. A high-quality draft genome of Oropetium was recently sequenced, but the lack of a chromosome scale assembly has hindered comparative analyses and downstream functional genomics. Here, we reassembled Oropetium, and anchored the genome into ten chromosomes using Hi-C based chromatin interactions. A combination of high-resolution RNAseq data and homology-based gene prediction identified thousands of new, conserved gene models that were absent from the V1 assembly. This includes thousands of new genes with high expression across a desiccation timecourse. The sorghum and Oropetium genomes have a surprising degree of chromosome-level collinearity, and several chromosome pairs have near perfect synteny. Other chromosomes are collinear in the gene rich chromosome arms but have experienced pericentric translocations. Together, these resources will be useful for the grass comparative genomic community and further establish Oropetium as a model resurrection plant.

## Introduction

Desiccation tolerance evolved as an adaptation to extreme and prolonged drying, and resurrection plants are among the most resilient plants on the planet. The molecular basis of desiccation tolerance is still largely unknown, but a number of models have emerged to dissect the genetic control of this trait (Hoekstra et al., 2001; Zhang and Bartels, 2018). The genomes of several model resurrection plants have been sequenced including *Boea hygrometrica* (Xiao et al., 2015), *Oropetium thomaeum* (VanBuren et al., 2015), *Xerophyta viscosa* (Costa et al., 2017), *Selaginella lepidophylla* (VanBuren et al., 2018), and *Selaginella tamariscina* (Xu et al., 2018). To date, no chromosome scale assembles are available for these species, limiting large-scale quantitative genetics and comparative genomics based approaches. Many resurrection plants are polyploidy or have prohibitively large genomes including those in the genera *Boea*, *Xerophyta*, *Eragostis*, *Sporobolus*, and *Craterostigma*. This complexity complicates genome assembly and gene redundancy in the polyploid species hinders downstream functional genomics work.

*Oropetium thomaeum* (hereon referred to as Oropetium), is a diploid resurrection plant and has the smallest genome among the grasses (245 Mb) (Bartels and Mattar, 2002). Oropetium plants are similar in size to Arabidopsis, but significantly smaller than the model grasses *Setaria italica* (Li and Brutnell, 2011) and *Brachypodium distachyon* (Brkljacic et al., 2011), with a short generation time of ~4 months. Oropetium is in the Chloridoideae subfamily of grasses and is closely related to the orphan cereal crops tef (*Eragrostis* tef) and finger millet (*Eleusine* coracana). Desiccation tolerance evolved independent several times within Chloridoideae (Gaff, 1977; Gaff and Latz, 1978; Gaff, 1987) making it a useful system for studying convergent evolution. Together, these traits make Oropetium an attractive model for exploring the origin and molecular basis of desiccation tolerance. Oropetium was one of the first plants to be sequenced using the long reads of PacBio technology, and the assembly quality was comparable to early Sanger sequencing based plant genomes such as rice and Arabidopsis (VanBuren et al., 2015). Despite the high contiguity of Oropetium V1, the assembly has 625 contigs and the BioNano based genome map was unable to produce chromosome-scale scaffolds. Furthermore, the V1 annotation was based on limited transcript evidence, and a high proportion of conserved plant genes were missing (VanBuren et al., 2015). Here, we reassembled the Oropetium genome using a more refined algorithm, and generated a chromosome scale assembly using Hi-C based chromatin interactions. The annotation quality was improved using high-resolution RNAseq data and protein homology, facilitating detailed comparative genomics with other grasses.

## Results

The first version of the Oropetium genome (V1) was sequenced with high coverage PacBio data (~72x) followed by error correction and assembly using the hierarchical genome assembly process (HGAP) (VanBuren et al., 2015). We reassembled this PacBio data using the Canu assembler (Koren et al., 2017a), which can more accurately assemble and phase complex repetitive regions. The resulting Canu based assembly (hereon referred to as V1.2) had fewer contigs than the V1 HGAP assembly, but had otherwise similar assembly metrics (Table 1).

**Table 1:**
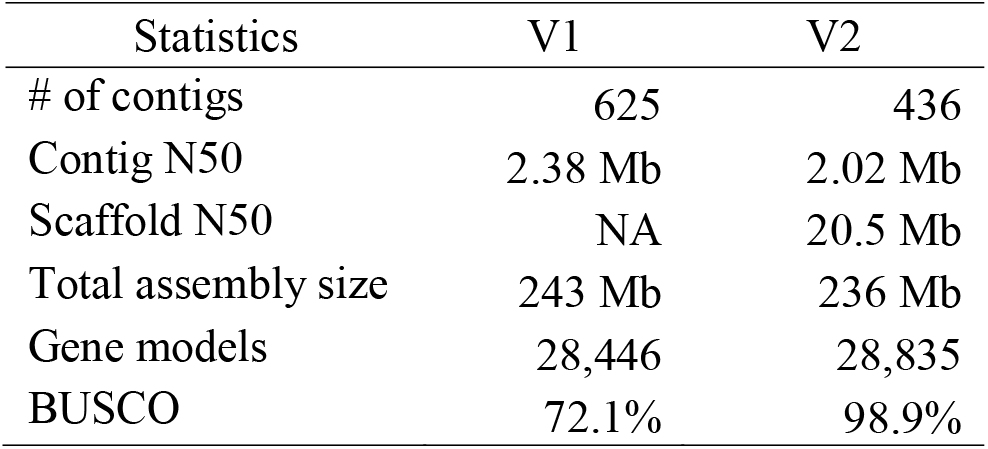
Comparison of the Oropetium V1 and V2 assembly and annotation statistics

Draft contigs were polished using a two-step process to remove residual insertion/deletion (indel) and single nucleotide errors. Contigs were first polished using the raw PacBio data with Quiver(Chin et al., 2013), followed by four rounds of reiterative polishing with Pilon (Walker et al., 2014) using high coverage Illumina paired end data. The final V1.2 assembly contains 436 contigs with an N50 of 2.0 Mb and total assembly size of 236 Mb. This is six megabases smaller than the V1 assembly, with slightly lower contiguity. More intact long terminal repeat retrotransposons (LTR-RTs) and centromere specific repeat arrays were identified in Oropetium V1.2 compared to V1, suggesting the Canu assembler resolved these repetitive elements more accurately. Thus, V1.2 was used for pseudomolecule construction.

The Oropetium V1.2 contigs were ordered and oriented into chromosome-scale pseudomolecules using high-throughput chromatin conformation capture (Hi-C). Hi-C leverages long-range interactions across distal regions of chromosomes to order and orient contigs. This approach is similar to genetic map-based anchoring, but with higher resolution. Illumina data generated from the Hi-C library was mapped to the V1.2 Oropetium genome using bwa (Li, 2013) and the proximity-based clustering matrix was generated using the Juicer and 3d-DNA pipelines (Durand et al., 2016; Dudchenko et al., 2017). After filtering and manual curation, ten high confidence clusters were identified (Figure 1). These ten clusters correspond to the haploid chromosome number of Oropetium. Regions with low density interactions highlight the centromeric and pericentromeric regions, and regions with higher than expected interactions represent topologically associated domains. After splitting six chimeric PacBio contigs, 239 contigs were anchored and oriented into ten chromosomes spanning 226.5 Mb or 95.8 % of the total assembled genome (Table 1). Chromosomes range in size from 11.0 to 34.7 Mb with an average size of 22.6 Mb. Most of the unanchored contigs are small (average size 42kb), or are entirely composed of rRNA, centromeric repeat arrays, or centromere specific LTR-RTs. Telomeres were identified at both ends of Chromosomes 1, 2, 3, 4, 5, 7, and 9 and on one end of Chromosomes 6, 8, and 10. Three unanchored contigs contain the remaining telomeres. This supports the completeness and accuracy of the pseudomolecule construction.

**Figure 1.**
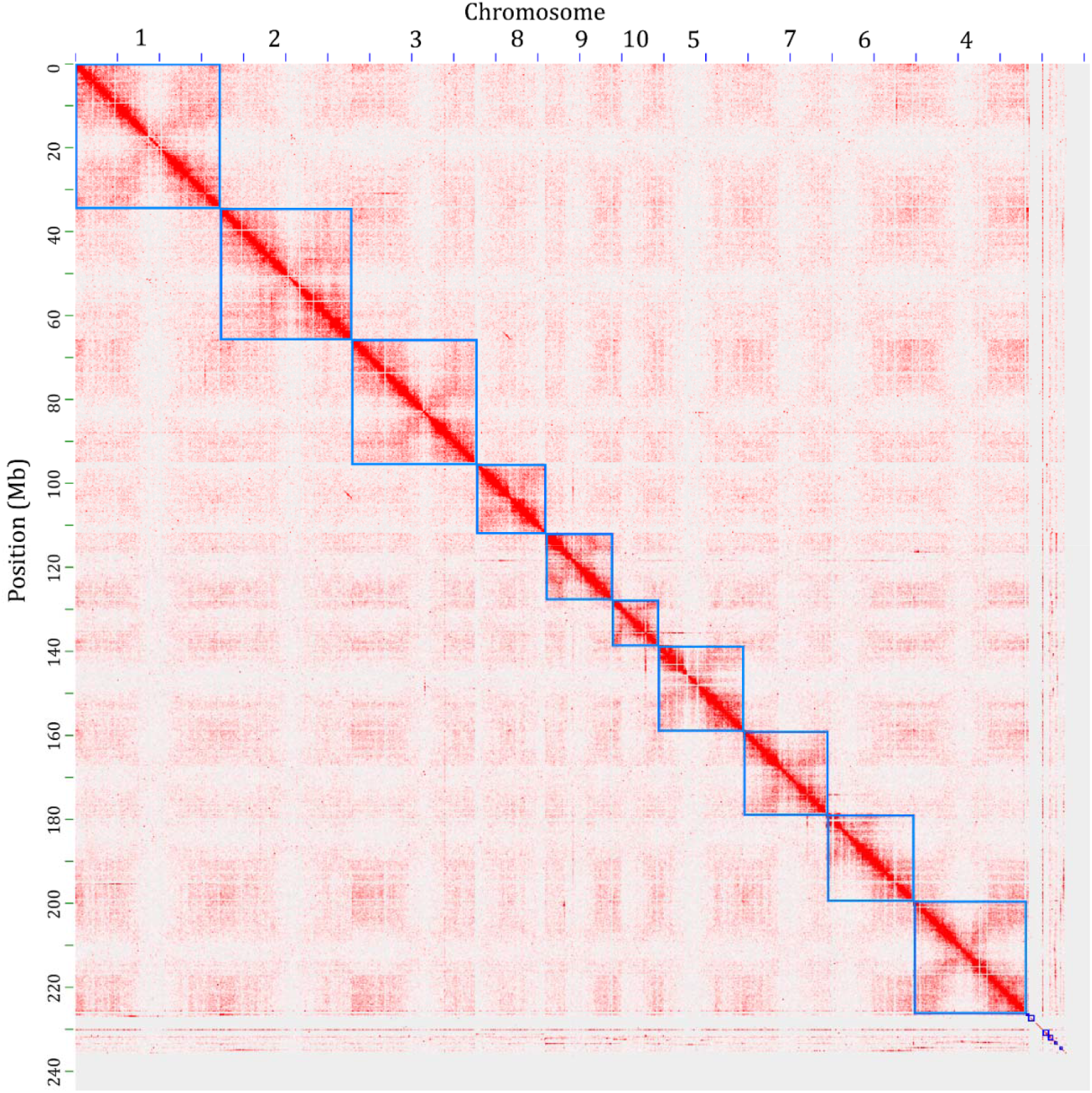
Hi-C based contig anchoring. Post-clustering heat map showing density of Hi-C interactions between contigs from the Juicer and 3d-DNA pipeline. The ten Oropetium chromosomes are highlighted by blue squares.

The chromosome scale Oropetium genome (hereon referred to as V2) was reannotated using the homology-based gene prediction program GeMoMa (Keilwagen et al., 2016; Keilwagen et al., 2018). Protein coding sequences from 11 angiosperm genomes and RNAseq data from Oropetium (VanBuren et al., 2017) were used as evidence. After filtering gene models derived from transposases, the final annotation consists of 28,835 high-confidence gene models. The annotation completeness was assessed using the Benchmarking Universal Single-Copy Ortholog (BUSCO) embryophyta dataset. The V2 gene models have a BUSCO score of 98.9%, suggesting the updated annotation is high-quality. In comparison, the Oropetium V1 annotation has a BUSCO score of 72%, and many conserved gene models were likely missing or mis-annotated. Nearly forty percent (11,227) of the gene models in V2 are new and were unannotated in V1. In addition, 10,837 gene models from V1 were removed or substantially improved in the V2 annotation. These discarded gene models either had little support based on protein homology to other species and transcript evidence from Oropetium, or they were misannotated transposable elements. In total, 94.3% of the gene models (27,216) were anchored to the ten chromosomes. Among the newly annotated gene models are 3,525 tandem gene duplicates (Figure 2a). Tandem duplicates span 3,062 arrays with 7,760 total genes. Of the arrays containing three or more genes, only 49 are new to V2, and the majority contain genes previously identified in V1. The boundaries of tandem duplicates are difficult to correctly annotate, resulting in fusions of two or more gene copies. The homology based annotation used in V2 was able to parse previously fused gene models.

**Figure 2.**
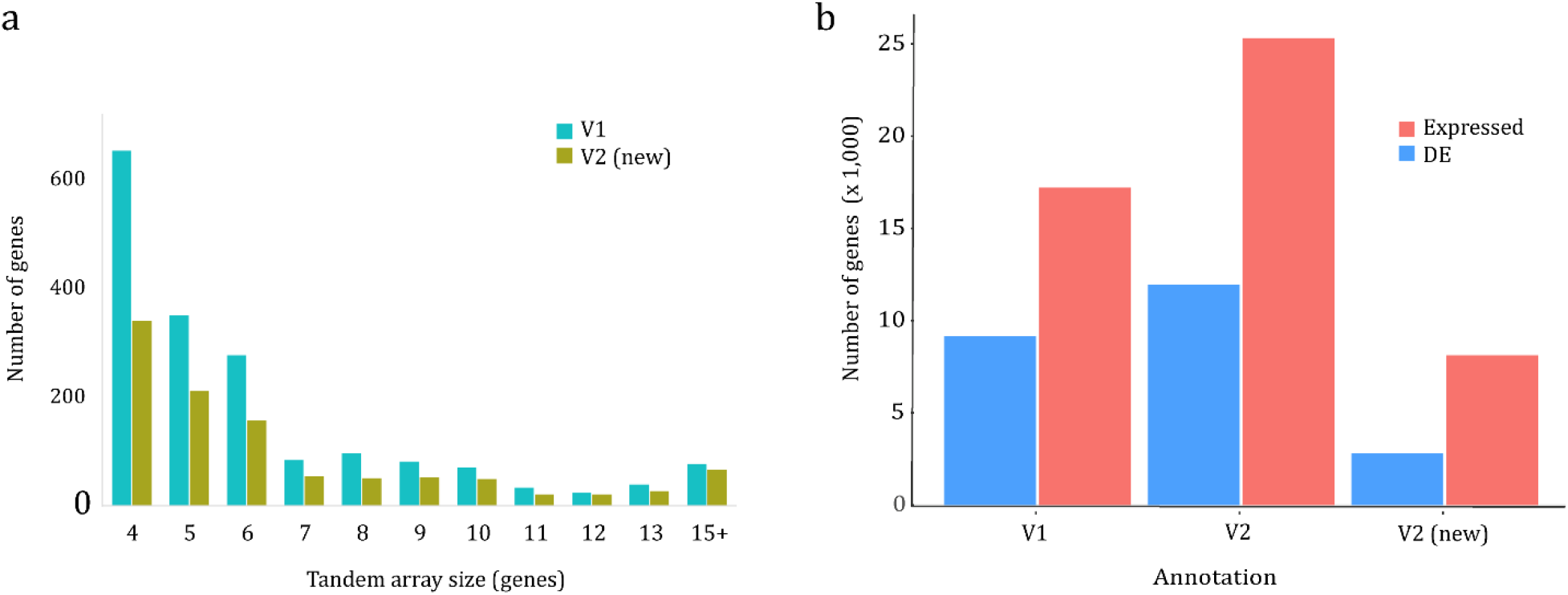
Characterization of the updated V2 Oropetium annotation. (a) Tandem gene array size comparison of the V1 and V2 annotation. Tandem genes identified in V1 are shown in blue and tandem genes newly annotated in V2 are shown in gold. (b) Comparison of expression patterns from the V1 and V2 annotation. The total number of genes with detectable expression and differential expression (DE) in the Oropetium desiccation/rehydration timecourse are plotted.

The expressions pattern of newly annotated genes was surveyed using previously generated RNAseq data (VanBuren et al., 2017). These timecourse datasets consist of seven samples from dehydrating and rehydrating Oropetium leaf tissue. Differentially expressed genes were identified based on comparisons of well-watered leaves with each dehydration and rehydration timepoint. In addition, each timepoint was compared with the timepoint immediately following it in the timecourse (ie. day 7 dehydration vs day 14). In the V1 annotation, 17,204 genes had detectable expression (count > 0 in at least one sample) compared to 25,314 genes in V2 (Figure 2b). Of the expressed genes, 9,149 V1 and 11,948 V2 were classified as differentially expressed in at least one of the comparisons. Most newly annotated genes (8,110) have detectable expression in at least one of the seven timepoints, and the majority are expressed in all timepoints. In total, 2,799 new V2 gene models were differentially expressed, suggesting the new genes have important and previously uncharacterized roles in desiccation tolerance.

We used the chromosome scale assembly of Oroeptium to survey patterns of genome organization and evolution related to maintaining a small genome size. The proportion of LTR-RTs in Oropetium V1 and V2 is similar, though V2 has more intact elements. LTR-RTs are the most abundant repetitive elements in Oropetium and collectively span 27% (62 Mb) of the genome. LTR-RTs are distributed non-randomly across the genome, and peaks of Gypsy LTR-RTs are observed in each of the ten chromosomes (Figure 3). These peaks of Gypsy LTR-RTs correspond to the pericentromeric regions. The pericentromeric regions show reduced intrachromosomal interactions in the Hi-C matrix, and contain arrays of centromeric repeats. The Oropetium V2 genome contains 8,965 155 bp monomeric centromeric repeats; considerably more than the 4,315 identified in the V1 assembly. The centromeric array sizes vary from 61 kb in chromosome 10 to 1,598 kb in Chromosome 4 (Figure 3; Table 2). Array sizes are likely underestimated, as only 52% of centromeric arrays were anchored to chromosomes, and 23 unanchored contigs contain centromeric repeat arrays. Gene density is low in the pericentromeric regions, consistent with the rice, Sorghum, Maize, and Brachypodium genomes (Paterson et al., 2009; Initiative, 2010; Du et al., 2017; Jiao et al., 2017). Collectively, pericentromeric regions span 67.5 Mb or 29% of the genome, a much smaller proportion than sorghum (62%; 460 Mb) (Paterson et al., 2009), but higher than rice (15%; 63 Mb) (Goff et al., 2002). The majority of intact LTRs (86%; 628) have an insertion time of less than one million years ago, with a steep drop off of insertion time after 0.4 MYA. This suggests LTRs are rapidly fragmented and purged in Oropetium to maintain its small genome size.

**Figure 3.**
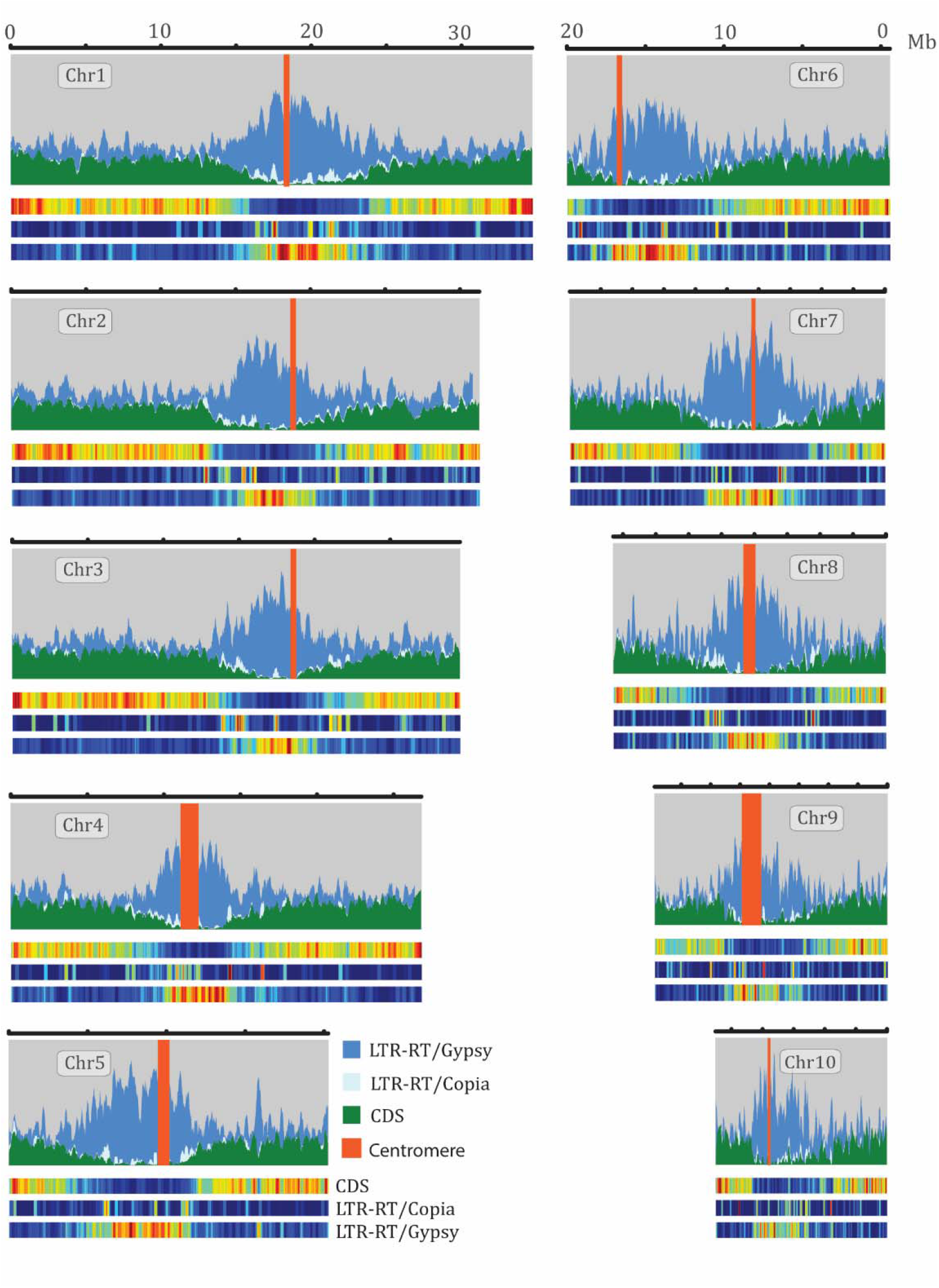
Landscape of the Oropetium genome. *Gypsy* and *Copia* long terminal repeat retrotransposons (LTR-RT) and CDS density are plotted for the ten Oropetium chromosomes. Features are plotted in sliding windows of 50kb with 25kb step size. The location of centromere specific tandem arrays is highlighted by red bars. The heatmaps below each landscape show relative density with red indicating high density and blue indicating low density for each feature.

**Table 2.** Centromeric repeat array composition

| **Chromosome** | **Start Cent.**<br>**Array (bp)** | **End Cent.**<br>**Array (bp)** | **Number of**<br>**Cent. Repeats** | **Cent. Size (bp)** |
| --- | --- | --- | --- | --- |
| Chr_1 | 18,899,082 | 19,114,162 | 154 | 215,080 |
| Chr_2 | 18,277,215 | 18,463,229 | 786 | 186,014 |
| Chr_3 | 18,882,303 | 18,993,598 | 308 | 111,295 |
| Chr_4 | 11,739,636 | 13,338,554 | 176 | 1,598,918 |
| Chr_5 | 10,361,368 | 10,828,355 | 800 | 466,987 |
| Chr_6 | 3,649,010 | 3,746,417 | 513 | 97,407 |
| Chr_7 | 12,434,273 | 12,559,564 | 272 | 125,291 |
| Chr_8 | 8,288,262 | 9,010,114 | 306 | 721,852 |
| Chr_9 | 6,142,739 | 7,433,209 | 1,044 | 1,290,470 |
| Chr_10 | 3,147,692 | 3,209,432 | 155 | 61,740 |
| Unanchored |  |  | 4,258 | 982,774 |

Previous comparative genomics analyses supported a high degree of collinearity between Oropetium and other grass genomes, but the draft assembly prevented detailed chromosome level comparisons. To date, no chromosome scale assemblies are available for other Chloridoideae grasses, though a draft genome is available for the orphan grain crop tef (*Eragrostis tef*) (Cannarozzi et al., 2014). We compared the V2 Oropetium chromosomes to the high-quality BTX 623 Sorghum genome (McCormick et al., 2018). Sorghum is in the Panicoideae subfamily of grasses which diverged from the ancestors of Chloridoideae ~31 MYA (Cotton et al., 2015). Despite this divergence, the ten chromosomes in Oropetium are largely collinear to the corresponding ten chromosomes in Sorghum, though large-scale inversions and translocations were identified (Figure 4a). Oropetium chromosomes 5, 6, and 8 are collinear along their length to sorghum chromosomes 9, 6, and 5 respectively. Oropetium chromosomes 1, 2, 4, and 7, are collinear to the arms of sorghum chromosomes 4, 10, 1, and 2, but the pericentric regions have translocated to other chromosomes. Oropetium chromosome 9 and sorghum chromosome 7 are syntenic but have two large-scale inversions, and Oropetium and sorghum chromosome 3 are syntenic with one inversion.

**Figure 4.**
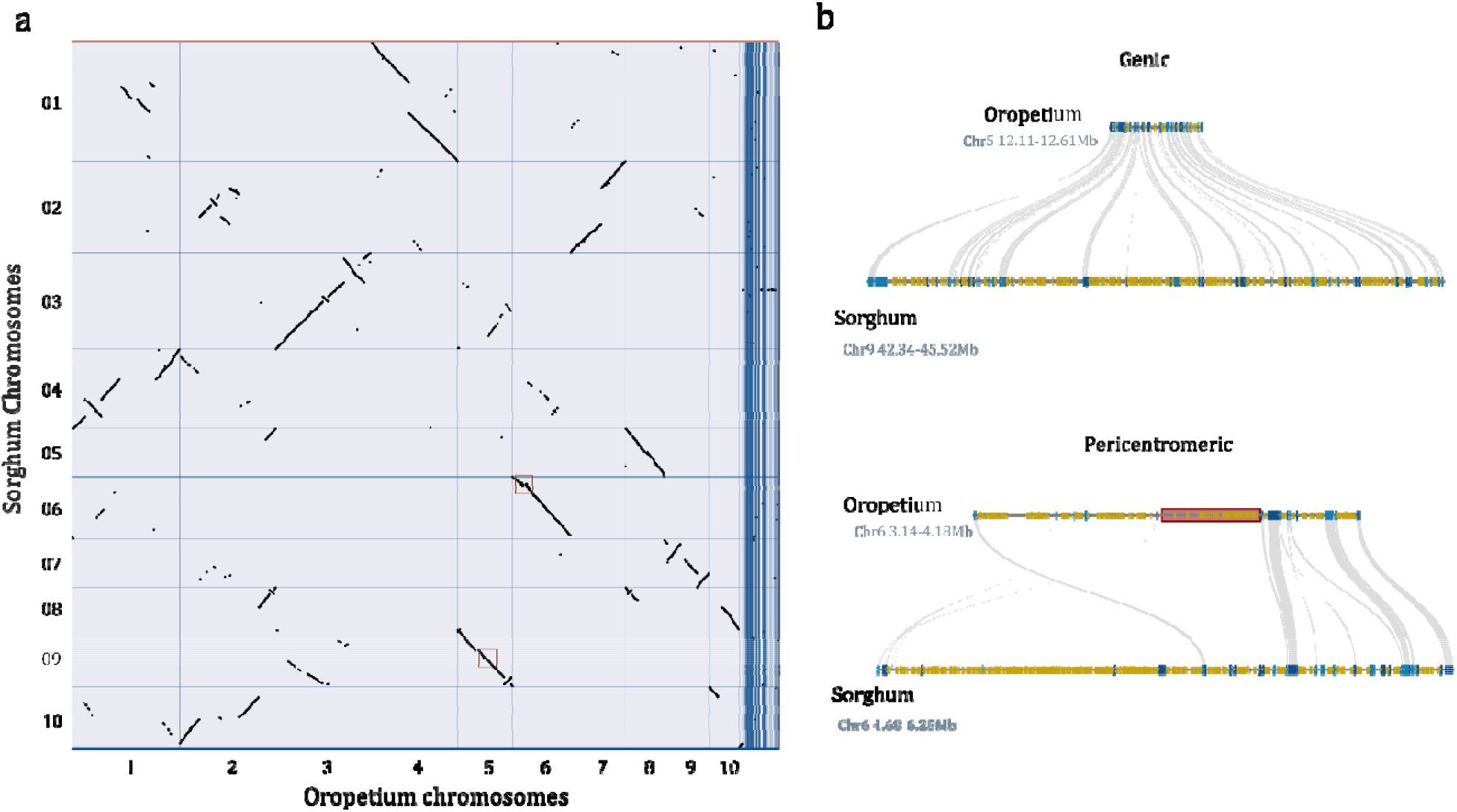
Comparative genomics between Oropetium and Sorghum. (a) Macrosyntenic dotplot of the Oropetium and Sorghum chromosomes based on 18,889 gene pairs. Each black dot represents a syntenic region between the two genomes. (b) Microsynteny of a typical genic region of Sorghum and Oropetium (top) and the pericentromeric region of Chromosome 6 of Oropetium and Sorghum (bottom). LTR-RTs are shown in yellow and genes are shown in blue. Syntenic orthologs are connected by gray lines. The centromeric repeat array in Oropetium is shown in red.

The sorghum genome is roughly three fold larger than Oropetium, and genome size dynamics in grasses are driven by purge and accumulation of retrotransposons (Wicker et al., 2010). Gene rich regions of Oropetium are 2-3x more compact than orthologous regions in sorghum, and much of this expansion in sorghum is caused by intergenic blocks of LTR-RTs (Figure 4b), consistent with patterns observed in the V1 assembly (VanBuren et al., 2015). The chromosome-scale nature of Oropetium V2 allowed us to survey patterns of collinearity in the pericentromeric regions. These regions have a lower degree of synteny with sorghum compared to gene rich euchromatin, consistent with retrotransposon-mediated rearrangements (Figure 4b). Pericentromeres are greatly expanded in Oropetium compared to the gene rich euchromatic blocks, similar to patterns observed in sorghum. The low gene density and low collinearity hinder detailed comparisons between pericentromeric regions.

## Discussion

The Oropetium V1 assembly quality rivals the early Sanger based genomes, and is much higher than the wealth of plant genomes assembled from short read Illumina sequences. Despite the high contiguity, the assembly was not chromosome scale, and essential genes were unannotated because of limited transcript evidence. This reflects the need to improve even the highest quality plant genomes. Our updated V2 Oropetium assembly better captures the gene space and allows for chromosome scale comparisons. The updated annotation includes thousands of new genes with differential expression related to desiccation tolerance. Hi-C based chromatin interactions anchored highly repetitive contigs across the pericentromeres, which are challenging to anchor using a classic genetic or optical map based approach. Together, these resources provide a useful outgroup for comparative genomics across the panicoid grasses and serve as a valuable foundation for functional genomics in this emerging model grass species.

## Methods

### Genome reassembly

The raw PacBio reads from the Oropetium V1 release (VanBuren et al., 2015) were reassembled with improved algorithms to better resolve highly complex and repetitive regions. PacBio data was error corrected and assembled using Canu (V1.4)(Koren et al., 2017b) with the following modifications: minReadLength=1500, GenomeSize=245Mb, minOverlapLength=1000. Other parameters were left as default. The resulting assembly graph was visualized in Bandage (Wick et al., 2015). The assembly graph was free of heterozygosity related bubbles, but many nodes (contigs) were interconnected by a high copy number retrotransposon. The Canu based contigs (assembly V1.2) were first polished using Quiver(Chin et al., 2013) with the raw PacBio data and default parameters. Contigs were further polished with Pilon (V1.22)(Walker et al., 2014) using ~120x coverage of paired-end 150 bp Illumina data. Quality-trimmed Illumina reads were aligned to the draft contigs using bowtie2 (V2.3.0) (Langmead and Salzberg, 2012) with default parameters. The overall alignment rate was 95.5%, which was slightly higher than alignment against the HGAP V1 assembly (94.5%). The following parameters for Pilon were modified: – flank 7, --K 49, and --mindepth 25. Other parameters were left as default. Pilon was run four times with an updated reference and realignment of Illumina data after each iteration. Indel corrections plateaued after the third iteration, suggesting polishing removed most residual assembly errors.

### HiC library construction analysis, and genome anchoring

Oropetium plants were maintained under day/night temperatures of 26 and 22°C respectively, with a light intensity of 200 μE m^−2^ sec^−1^ and 16/8 hr photoperiod. Young leaf tissue was used for HiC library construction with the Proximo™ Hi-C Plant kit (Phase Genomics) following the manufactures protocol. Briefly, 0.2 grams of fresh, young leaf tissue was finely chopped and the chromatin was immediately crosslinked. The chromatin was fragmented and proximity ligated, followed by library construction. The final library was size selected for 300-600 bp and sequenced on the Illumina HiSeq 4000 under paired-end 150 bp mode. Adapters were trimmed and low-quality sequences were removed using Trimmomatic (V0.36) (Bolger et al., 2014). Read pairs were aligned to the Oropetium contigs using bwa (V0.7.16)(Li, 2013) with strict parameters (-n 0) to prevent mismatches and non-specific alignments in duplicated and repetitive regions. SAM files from bwa were used as input in the Juicer pipeline, and PCR duplicates with the same genome coordinates were filtered prior to constructing the interaction based distance matrix. In total, 101 filtered read pairs were used as input for the Juicer and 3d-DNA HiC analysis and scaffolding pipelines (Durand et al., 2016; Dudchenko et al., 2017). Contig ordering, orientation, and chimera splitting was done using the 3d-DNA pipeline(Dudchenko et al., 2017) under default parameters. Contig misassemblies and scaffold misjoins were manually detected and corrected based on interaction densities from visualization in Juicebox. In total, six chimeric contigs were identified and split at the junction with closest interaction data. The manually validated assembly was used as input to build the ten scaffolds (chromosomes) using the finalize-output.sh script from 3d-DNA. Chromosomes and unanchored contigs were renamed by size, producing the V2 assembly.

### Genome annotation

The Oropetum V2 assembly was reannotated using the homology-based gene prediction program Gene Model Mapper (GeMoMa: V 1.5.2) (Keilwagen et al., 2016; Keilwagen et al., 2018). GeMoMa uses protein homology and RNAseq evidence to predict gene models. Genome assemblies and gene annotation for the following 11 species were downloaded from Phytozome (V12) and used as homology based evidence: *Arabidopsis thaliana*, *Brachypodium distachyon*, *Glycine max*, *Oryza sativa*, *Panicum hallii*, *Populus trichocarpa*, *Prunus persica*, *Setaria italica*, *Solanum lycopersicum*, *Sorghum bicolor*, *Theobroma cacao*. Translated coding exons and proteins from the reference gene annotations and genome assemblies were extracted using the module Extractor function of GeMoMa (module Extractor: Ambiguity=AMBIGUOUS, r=true). RNAseq data from Oropetium desiccation and rehydration timecourses (VanBuren et al., 2017) was aligned to the V2 Oropetium genome using HISAT2 (Kim et al., 2015) with default parameters. The resulting BAM files were used to extract intron and exon boundaries using the module ERE (module ERE: s=FR_FIRST_STRAND, c=true). translated coding exons from other species were aligned to the Oropetium genome using tblastn and transcripts were predicted based on each reference species independently using the extracted introns and coverage (module GeMoMa). Finally, the predictions based on the 11 reference species were combined to obtain a final prediction using the module GAF. Gene models containing transposases were filtered, resulting in a final annotation of 28,835 gene models. The annotation completeness was assessed using the plant specific Benchmarking Universal Single-Copy Ortholog (BUSCO) dataset (version 3.0.2, embryophyta_odb9) (Simão et al., 2015). The following report was obtained from BUSCO: 98.9% overall, 95.4% single copy, 3.5% duplicated, 0.6% fragmented, 0.5% missing. Gene model names from V1 were conserved where possible, and new gene models received new names.

### Expression analysis

Oropetium RNAseq data from desiccation and rehydration timecourses was reanalyzed using the updated gene model annotations (VanBuren et al., 2017). Four time points during dehydration (days 7, 14, 21, and 30), two during rehydration (24 and 48 hours), and one well-watered sample were analyzed. Based on principle component analysis, replicate 2 of the ‘well-watered and ‘D21’ samples were excluded from the analysis. Each other timepoint had three replicates. Gene expression was quantified on a transcript level using salmon (v 0.9.1) in quasi-mapping mode (Patro et al., 2017). Default parameters were used with the internal GC bias correction in salmon. The R package tximport (v 1.2.0) was used to map transcript level quantifications to gene level counts (Team, 2013; Soneson et al., 2015). We conducted differential expression analysis with the remaining samples using the R package DESeq2 (v 1.14.1) set to default parameters [3,4].

### Identification of LTR-RTs

A preliminary list of candidate long terminal repeat retrotransposons (LTR-RTs) from Oropetium were identified using LTR_Finder (V1.02) (Xu and Wang, 2007) and LTRharvest (Ellinghaus et al., 2008). The following parameters for LTRharvest were modified: –similar 90 – vic 10 –seed 20 –minlenltr 100 –maxlenltr 7000 –mintsd 4 –maxtsd 6 –motif TGCA –motifmis 1. LTR_Finder parameters were: -D 15000 –d 1000 –L 7000 –1 100 –p 20 –C –M 0.9. LTR_retriever(Ou and Jiang, 2017) was used to filter out false LTR retrotransposons using the target site duplications, terminal motifs, and Pfam domains. Default parameters were used for LTRretriever. LTRretirever produced a list of full length, high-quality LTRs. LTRs were annotated across the genome using RepeatMasker (http://www.repeatmasker.org/)(Smit et al., 1996) and the non-redundant LTR-RT library constructed by LTR_retriever. The insertion time of intact LTRs was calculated in LTR_retriever using the formula T=K/2μ with a neutral mutation rate of μ=1 × 10-8 mutations per bp per year.

### Comparative genomics

Syntenic gene pairs between the Oropetium and Sorghum genomes were identified using the MCSCAN toolkit (V1.1) (Wang et al., 2012) implemented in python (https://github.com/tanghaibao/jcvi/wiki/MCscan-(Python-version)). Default parameters were used. Gene models were aligned using LAST and hits were filtered to find the best 1:1 syntenic blocks. Macrosyntenic dotplots were constructed in MCScan.

## Availability of supporting data

The V2 Oropetium genome assembly and updated annotation can be downloaded from CoGe (https://genomeevolution.org/coge) under Genome ID 51527 and from Phytozome (https://phytozome.jgi.doe.gov/pz/portal.html). The raw Hi-C Illumina data has been deposited on the Short Read Archive (SRA) under NCBI BioProject ID PRJNA481965.

